# Lipin-1 regulates lipid catabolism in pro-resolving macrophages

**DOI:** 10.1101/2020.06.03.121293

**Authors:** Robert M. Schilke, Cassidy M.R. Blackburn, Shashanka Rao, David Krzywanski, Brian N. Finck, Matthew D. Woolard

**Affiliations:** Department of Microbiology and Immunology, Louisiana State University Health Sciences Center, Shreveport, LA 71130; Department of Cellular Biology and Anatomy, Louisiana State University Health Science Center, Shreveport, LA 71130; Division of Geriatrics and Nutritional Science, Washington University School of Medicine, St. Louis, USA

**Keywords:** Lipin-1, macrophage, mitochondria, metabolism, efferocytosis

## Abstract

Macrophages reprogram their metabolism to promote appropriate responses. Pro-resolving macrophages primarily utilize fatty acid oxidation as an energy source. Metabolites generated during the catabolism of fatty acids aid in the resolution of inflammation and tissue repair, but the regulatory mechanisms that control lipid metabolism in macrophages is not fully elucidated. In this current study we show that lipin-1, a phosphatidic acid phosphatase and regulator of lipid metabolism, is required for increased oxidative phosphorylation during IL-4 mediated responses. We also show that the transcriptional coregulatory function of lipin-1 is required for β-oxidation in response to palmitate (free fatty acid) and apoptotic cell derived lipids. BMDMs lacking lipin-1 have a reduction in critical TCA cycle metabolites following IL-4 stimulation, suggesting a break in the TCA cycle that is supportive of lipid synthesis rather than lipid catabolism. Together, our data demonstrate that lipin-1 regulates intermediary metabolism within pro-resolving macrophages and highlights the importance of aligning macrophage metabolism with proper responses to stimuli.

## Introduction

Macrophages are innate immune cells that regulate tissue homeostasis and are critical for disease resolution. Macrophages are able to polarize toward distinct phenotypes, such as pro-resolving macrophages (M2) macrophages that aid in wound healing processes[1, 2]. As macrophages polarize, they change their metabolic profile to effectively respond to stimuli[3]. Pro-inflammatory macrophages predominately rely on glycolysis to generate ATP. Oxidative phosphorylation is reduced to generate reactive oxygen species and promote lipid synthesis[4]. Pro-inflammatory macrophages have a break in the TCA cycle at two distinct points. The first break leads to the accumulation of citrate, which is exported out of the mitochondria to be used in lipid synthetic pathways. The second break leads to the accumulation of succinate which stabilizes HIF1α to increase expression of pro-inflammatory genes. Pro-inflammatory macrophages also increase the pentose phosphate pathway (PPP) to generates NADPH, which is used in lipid synthesis and cellular redox potential[5]. Pro-resolving macrophages initially increase glycolysis to quickly generate ATP, but heavily rely on oxidative metabolism to produce sufficient energy to carry out their functions. Pro-resolving macrophages utilize β-oxidation of fatty acids to produce acetyl-CoA, which is oxidized via a series of reaction in the TCA cycle. The oxidation of acetyl-CoA generates reducing equivalents, such as NADH, to fuel the electron transport chain[5]. B-oxidation has been shown to promote macrophage polarization towards the pro-resolving phenotype, and is essential to disease resolution [6]. Catabolism of apoptotic cell derived lipids promotes IL-10 production and subsequent wound healing in a mouse model of myocardial infarction[6]. However, how macrophages regulate lipid metabolism to promote pro-resolving functions is not fully understood.

Lipin-1 is a phosphatidic acid phosphatase that converts phosphatidic acid into diacylglycerol to be used in lipid synthetic pathways[7]. Lipin-1 is also a transcriptional coregulator that binds to transcription factors, such as PPARs, to regulate gene expression[8]. Lipin-1 has been shown to regulate the expression of genes that encode proteins involved in fatty acid transport and lipid catabolism in a variety of tissues[9]. Hepatic lipin-1 amplifies PGC-1α/PPARα regulatory pathway to stimulate β-oxidation[8]. Lipin-1 not only controls fatty acid metabolism, but may also regulate oxidative phosphorylation, since genetic depletion of lipin-1 in skeletal muscle results in a significant decrease in oxidative phosphorylation[10]. Loss of hepatic lipin-1 in mice results in an alteration in major metabolic pathways in a diurnal cycle, such as an increase in fatty acid synthesis and elevated peripheral glucose utilization [11]. Genetic depletion of lipin-1 in *C. elegans* results in an increase is fatty acid synthesis genes, with a corresponding decrease in lipolytic genes [12]. Individuals lacking lipin-1 have impaired β-oxidation during exercise, which can result in rhabdomyolysis [13]. Collectively, these studies demonstrate the contribution of lipin-1 to lipid catabolism.

Macrophages balance their cellular metabolism to carry out their function, and alterations in metabolic reprogramming prevent appropriate responses. The inability to breakdown lipid can be detrimental to macrophage function [1, 6, 14]. The two molecular function of lipin-1 seem to play contrasting roles in modulating macrophage polarization. Specifically, lipin-1 enzymatic activity promotes pro-inflammatory macrophage responses, while the nuclear activity has pro-resolving properties [7, 15, 16]. Lipin-1 mediated diacylglycerol production promotes the activation of the AP-1 transcription factor, which leads to the production of pro-inflammatory cytokines; IL-6, TNF-α and IL-12 and pro-inflammatory eicosanoids such as PGE_2_ [15, 16]. The enzymatic activity of lipin-1 in macrophages contributes to the pathogenesis of atherosclerosis, colitis, colon cancer, LPS-induce inflammation, and alcoholic liver disease[7, 16–18]. We have further demonstrated that lipin-1 can also contribute to pro-resolving macrophage polarization[19]. Specifically, our studies suggest that lipin-1 transcriptional coregulatory activity is required for promotion of macrophage wound healing phenotype. As evident by the fact that mice lacking lipin-1 from myeloid cells have accelerated atherosclerosis and a defect in wound healing, which we did not observe in mice lacking myeloid-associated lipin-1 enzymatic activity [19, 20]. However, the molecular mechanisms by which lipin-1 transcriptional co-regulatory activity promotes macrophages pro-resolving phenotypic activity is unknown. The contribution of lipin-1 to β-oxidation in other tissues, taken together with lipin-1 involvement in polarization of macrophages toward a pro-resolving phenotype led us to hypothesize that lipin-1 regulates β-oxidation in pro-resolving macrophages.

We used a mouse model to genetically deplete either only the enzymatic activity of lipin-1, or completely deplete lipin-1 within myeloid derived cells. This allows us to elucidate which activity of lipin-1 contributes to lipid metabolism within macrophages. We show that lipin-1 is required for increased oxidative metabolism in response to IL-4 as well as exogenous lipid in pro-resolving macrophages.

## Methods

### Animals

All animal studies were approved by the LSU Health Sciences Center-Shreveport institutional animal care and use committee. All animals were cared for according to the National Institute of Health guidelines for the care and use of laboratory animals.

All animals used in this study were 8 to 10-week old mice. Mice lacking lipin-1 enzymatic activity from myeloid cells (lipin-1^mEnzy^KO) were generated as previously reported [9]. Briefly, mice with exons 3 and 4 of the *Lpin1* gene flanked by LoxP sites (genetic background: C57BL/6J and SV129; generously provided by Brian Finck and Roman Chrast) were crossed with C57BL/6J LysM-Cre transgenic mice purchased from Jackson Laboratory (Bar Harbor, ME). Mice fully lacking lipin-1 from myeloid cells (lipin-1^m^KO) were generated by crossing mice with exon 7 of the *Lpin1* gene flanked by LoxP sites (genetic background: C57BL/6J and SV129; generously provided by Brian Finck [15]) with C57BL/6J LysM-Cre transgenic mice purchased from Jackson Laboratory (Bar Harbor, ME). Age matched lipin-1flox/flox littermate mice were used as controls.

### Generation of bone marrow-derived macrophages

Bone marrow-derived macrophages (BMDMs) were generated from lipin-1^mEnzy^KO, lipin-1^m^KO and littermate control mice as previously described[21]. Briefly, femurs were excised under sterile conditions and flushed with Dulbecco’s modified Eagle’s Knock out medium (DMEM; Gibco, 10829) supplemented with 5% fetal bovine serum (Atlanta biologicals, S11150), 2mM L-glutamax (Gibco - 35050-061), 100 U/ml penicillin-streptomycin (Cellgro, 30-604-CI), 1mM sodium pyruvate (Cellgro, 25-060-CI), and 0.2 % sodium bicarbonate (Quality biological, 118-085-721). Red blood cells were lysed using ammonium chloride-potassium carbonate (0.15 M NH_4_Cl, 10 mM KHCO_3_, 0.1 mM NA_2_EDTA, adjusted to pH 7.2 and filter sterilized in 0.22 μm filter) lysis (ACK) followed by PBS wash. Isolated cells were incubated in sterile petri dishes for 7 to 10 days in BMDM differentiation medium – DMEM (Gibco, 11965-092) supplemented with 30% L-cell conditioned medium, 20% fetal bovine serum (Atlanta biologicals, S11150) at 37°C and 5% CO2. Once cells were 80% confluent, they were collected using 10 mM EDTA, pH 7.6 treatment. BMDMs were resuspended in D-10 (DMEM (Gibco: 11965-092), 10% fetal bovine serum (Atlanta biologicals S11150), 2mM glutamax (Thermofischer 35050-061), 100U/mL penicillin-streptomycin (Cellgro), and 1mM sodium pyruvate (Cellgro) for further experiments

### Apoptotic cell (AC) generation

Thymocytes (Jurkats) were subjected to UV for 20 minutes in a 10cm dish in sterile PBS. Cells were collected and media was changed to D-10 complete media followed by a 2-hour incubation at 37°C to allow for time for cells to become apoptotic. Cells were collected, centrifuged at 400xg for 5 minutes. Cells were resuspended in 2×10^−6^ M PKH26 for 3 minutes. Staining was stopped by adding equivalent volumes of FBS.

### Mitochondrial Bioenergetics

BMDM collected from lipin-1^mEnzy^KO, lipin-1^m^KO and littermate control mice were seeded at a density of 150,00 cells/well on XF24 cell culture microplates and allowed to incubate for 4 hours. Experiments were conducted in XF assay medium containing 25 mM glucose, 2 mM L-glutamine, and 1 mM Na pyruvate, and analyzed using a Seahorse XF24 extracellular flux analyzer (Agilent Technologies). Where indicated, the following were injected: ATP-synthesis inhibitor oligomycin (1 μM), carbonyl cyanide 4-(trifluoromethoxy) phenylhydrazone (FCCP; 2 μM) to uncouple ATP synthesis, rotenone (100 nM) to block complex I, and antimycin A (1 μM) (Sigma) to block complex III. Oxygen consumption rate was analyzed using Wave Desktop software (Agilent Technologies).

### Palmitate utilization

BMDM collected from lipin-1^mEnzy^KO, lipin-1^m^KO, and littermate control mice were seeded at a density of 150,000 cells/well on XF24 cell culture microplates and allowed to incubate for 4 hours. Macrophages were treated with 40ng/mL of IL-4 for 1.5 hours. Media was then changed to nutrient restricted media containing Agilent base media (100840-000) supplemented with 0.5mM pyruvate (Cell gro, 25-060-CI) and 0.5mM glutamax (Gibco -35050-061). Cells were then pre-treated with 2mM 2-deoxy-D-glucose (2-DG) (Sigma-D6134) for 15 minutes prior to the assay. Directly before analysis, 100 μM palmitate was added and OCR was assayed via the Seahorse extracellular flux analyzer (Agilent).

### MerTK receptor staining

Zymosan (0.1mg) was injected intraperitoneally into lipin-1^m^KO and littermate control mice and allowed to incubate for 6 days. Mice were then sacrificed and the peritoneal lavage was collected in FACS wash buffer (1% bovine serum albumin, 1 mM EDTA, and 0.1% sodium azide in phosphate buffered saline) and incubated with CD16/CD32 (e-Bioscience, 14-0161-86) for 20 min. Cells were incubated with PECy7 conjugated CD11b (e-Biosciences, 25-0112-81,clone M1/70), and PE conjugated anti-MerTK antibody (Biolegend, 151506) for 30 minutes in the dark. Cells were spun at 400xg for 5 minutes and resuspended in FACS fix (1% PFA in FACS wash buffer). Appropriate F Minus One Controls were used to correct background and exclude spectral overlap staining. Compensation control (Comp Bead, Invitrogen, 01-2222-42) were used. Flow cytometry was performed using Quanteon flow cytometer (Acea Biosciences). Data analysis was performed using FCS express (Denovo Software) and NovoExpress (Acea Biosciences).

### Central carbon analysis

One mL of BMDM from lipin-1^m^KO and littermate control mice at a concentration of 1×10^6^/mL was placed into falcon tubes (VWR-60818-500). Cells were either treated with or without IL-4 (40ng/mL) for 4 hours. Cells were then collected and spun at 400g for 5 minutes. Cell pellets were sent to Creative proteomics (Shirly, NY) for central carbon analysis.

### Lipid uptake

500 uL of BMDM from lipin-1^mEnzy^KO, lipin-1^m^KO, and littermate control mice at a concentration of 1×10^6^/mL in HBSS (Fisher Scientific-SH30588.01) containing 25 mM glucose was placed into falcon tubes (VWR-60818-500). BODIPY labeled palmitate (Thermofisher-D3821) (100 μM) was added for macrophages and allowed to incubate for 1 hour. Cells were then washed and suspended in FACS Wash and MFI was analyzed via flow cytometry.

### In vivo efferocytosis

Lipin-1^mEnzy^KO, lipin-1^m^KO and littermate control mice were injected intraperitoneally with 0.1mg of zymosan. Thymocytes (Jurkat) were subjected to UV for 20 minutes followed by 2-hour incubation at 37°C to allow apoptosis to occur. Jurkat cells were then labeled with PKH-26. 6 days post zymosan injection, 4 million labeled AC were injected into the peritoneal cavity of the mice and allowed to incubate for 45 minutes. Mice were then sacrifice and peritoneal lavages were collected and stained with PECy7 conjugated anti-CD11b (e-Biosciences, 25-0112-81,clone M1/70), AF700 conjugated anti-CD45.2 (Biolegend,109821,clone104), FITC conjugated anti-Ly6G (BD Biosciences,551460, clone1A8), and APC conjugated anti-F4/80 (Invitrogen, 15-4801-80, clone BM8). Lavages were then analyzed via flow cytometry.

### Statistical analysis

GraphPad Prism 5.0 (La Jolla, CA) was used for statistical analyses. A student T Test analysis was used for comparison between two data sets. All other statistical significance was determined using a one-way ANOVA with a Dunnett’s post-test. All *in vivo* and *in vitro* experiments were performed a minimum of three times. Figure legends provide specific details for each data set.

## RESULTS

### Lipin-1 is required for increased metabolic activity during lipid catabolic states

Lipin-1 enzymatic activity promotes macrophage pro-inflammatory responses [15, 16], and we have recently demonstrated that lipin-1 transcriptional coregulatory activity contributes to IL-4 elicited gene expression in macrophages [19]. Interleukin-4 (IL-4) stimulation polarizes macrophages towards a pro-resolving macrophage, which heavily relies on β-oxidation and oxidative phosphorylation [5]. In hepatocytes and skeletal muscle cells, lipin-1 regulates β-oxidation and oxidative phosphorylation[8, 22]. We wanted to investigate if lipin-1 regulated oxidative metabolism in macrophages. We used BMDMs from lipin-1^mEnzy^KO (EKO), lipin-1^m^KO (KO), and littermate controls (WT) to determine the contribution of the enzymatic activity and infer the contribution of the transcriptional coregulatory activity towards oxidative metabolism. BMDMs from EKO, KO, and WT mice were stimulated with 40ng/ml of IL-4 for 4 hours and cellular bioenergetics were analyzed by Seahorse extracellular flux analyzer (Agilent Technologies). There were no differences in baseline respiration between WT and EKO macrophages (Figure 1A & 1B). IL-4 stimulation of both WT and EKO BMDMs resulted in a significant increase in oxygen consumption rate (OCR) (Figure 1C & 1D). This suggests that the enzymatic activity of lipin-1 is dispensable for IL-4 elicited oxygen consumption. We then wanted to test if lipin-1 transcriptional coregulatory activity contributes to oxidative metabolism in macrophages. WT and lipin-1 KO macrophages were stimulated with IL-4 for 4 hours and cellular bioenergetics were analyzed. There was no difference in baseline respiration between WT and lipin-1 KO BMDMs (Figure 2A & 2B). Lipin-1 KO BMDM failed to increase OCR in response to IL-4 stimulation, as seen in the WT BMDMs (Figure 2C & 2D). The inability of lipin-1 deficient macrophages to increase oxygen consumption rate in response to IL-4 suggests that lipin-1 transcriptional coregulator function is required for oxidative phosphorylation within macrophages.

**Figure 1:**
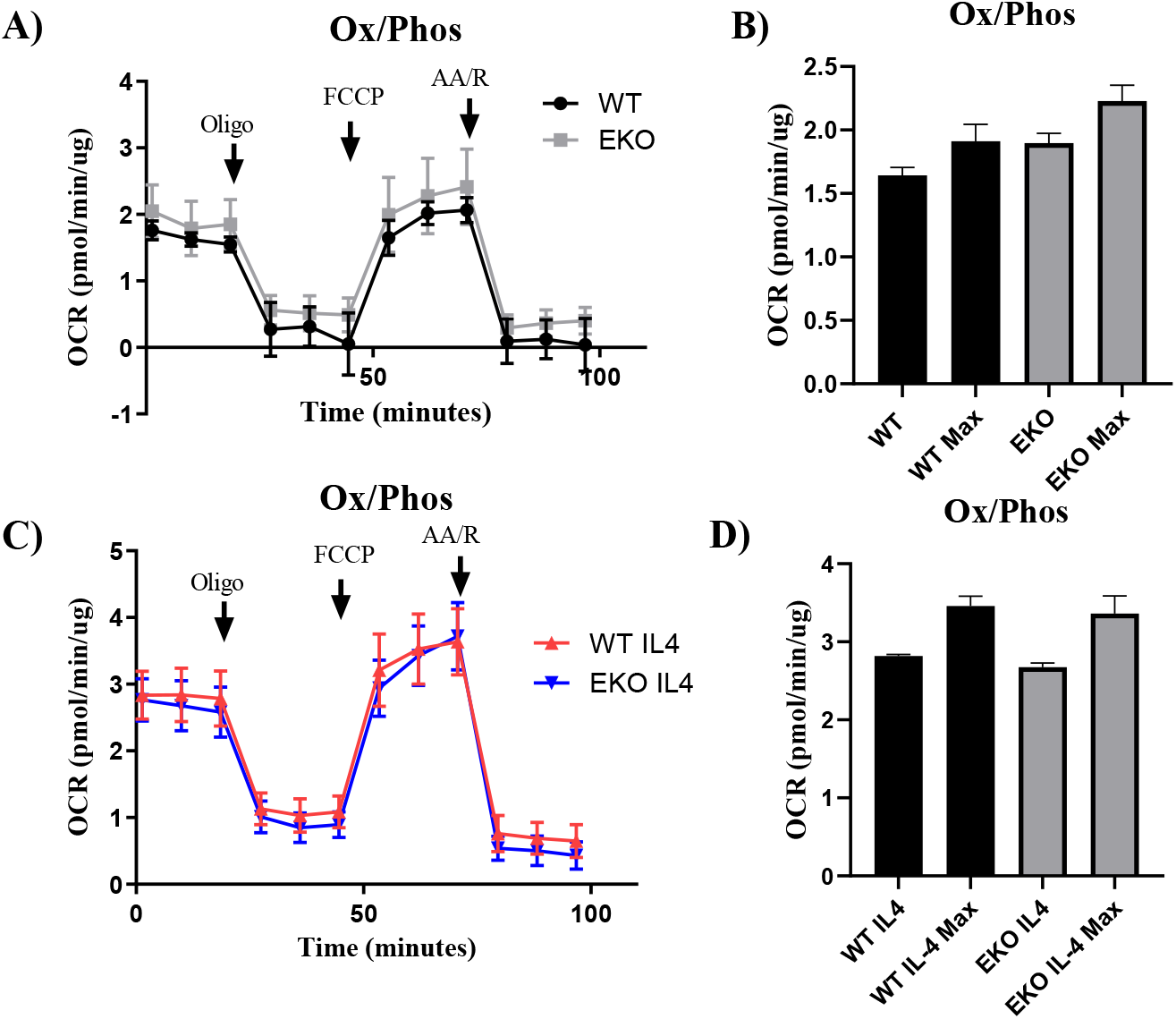
Lipin-1 enzymatic activity does not contribute to oxidative phosphorylation. BMDM isolated from WT and EKO macrophages were treated with or without 40ng/mL IL-4 for 4 hours. Oxygen consumption was analyzed via Seahorse extracellular flux analyzer. Arrows indicate injection of oligomycin (Oligo), FCCP, antimycin A and rotenone (AA/R). A) Untreated WT and EKO macrophages. B) Basal and maximal (FCCP) OCR of WT and EKO BMDM’s. C) WT and EKO BMDM’s treated with IL-4.

**Figure 2:**
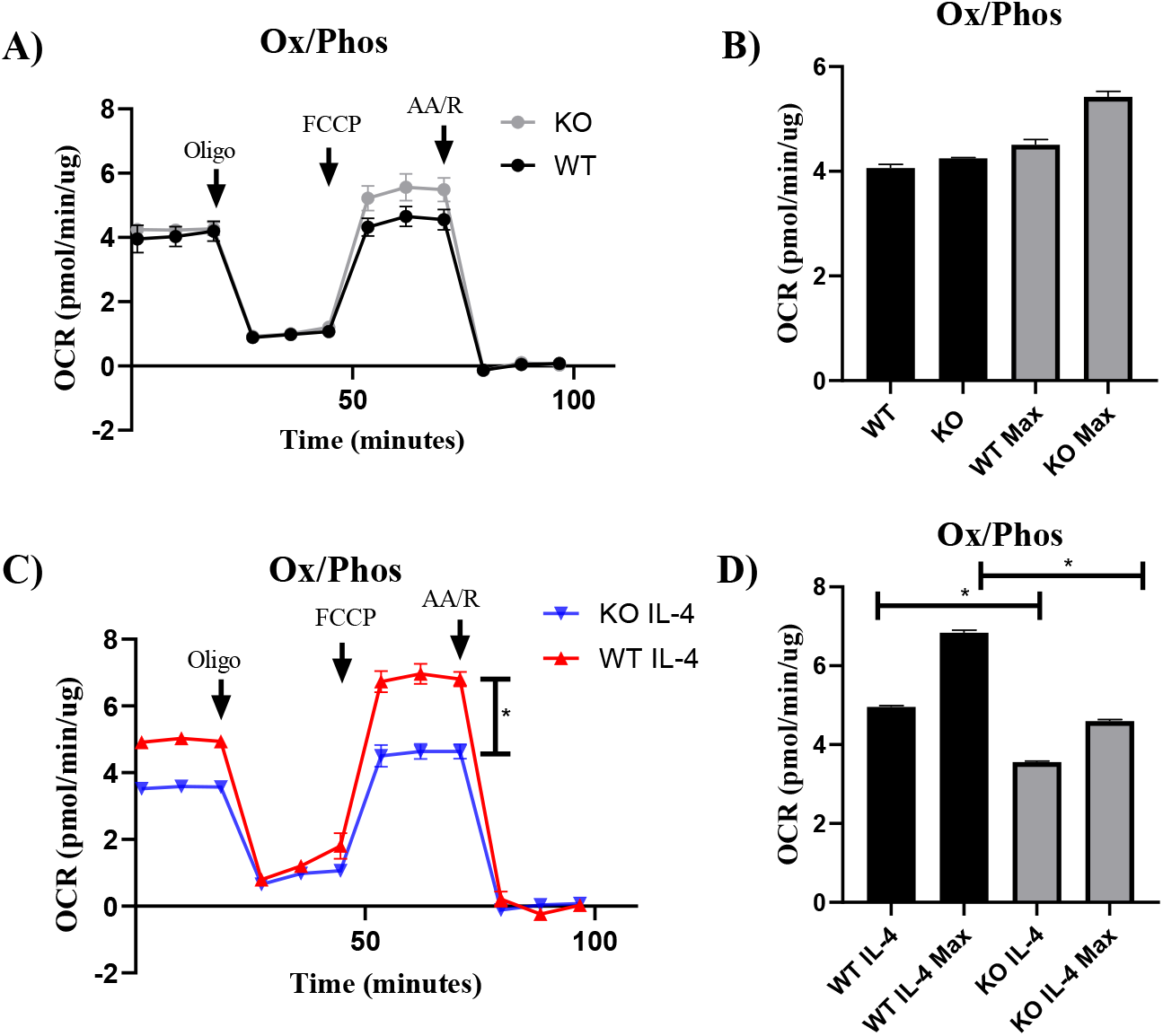
Lipin-1 transcriptional coregulator function contributes to oxidative phosphorylation. BMDM isolated from WT and KO macrophages were treated with or without 40ng/mL IL-4 for 4 hours. Oxygen consumption was analyzed via Seahorse extracellular flux analyzer. A) Untreated WT and KO macrophages. B) Basal and maximal (FCCP) OCR of WT and KO BMDM’s. C) WT and KO BMDM’s treated with IL-4. D) Basal and maximal (FCCP) OCR of WT and KO BMDM’s. N=3. Stars represent p<0.5.

### Lipin-1 regulates lipid utilization within macrophages

IL-4 stimulated macrophages shuttle fatty acids into the mitochondria to undergo β-oxidation and fuel oxidative phosphorylation [23]. The loss of lipin-1 resulted in a failure of macrophages to increase oxygen consumption in responses to IL-4, and we hypothesize this defect is due to an inability to breakdown lipids for energy leading to oxygen consumption. To determine if lipin-1 deficient macrophages can use lipids to promote oxygen consumption we created an environment to most favorable to using lipids for beta-oxidation. We stimulated WT and lipin-1 KO BMDM with IL-4 for 1.5 hours to elicit β-oxidation in nutrient (pyruvate and glutamine) restricted media. We inhibited glycolysis with the addition of 2DG and added 100-μM palmitate to promote lipid mediated oxygen consumption. OCR was measured via seahorse XFE analyzer. We observed no difference in oxygen consumption between IL-4 treated WT and lipin-1 KO macrophages not treated with palmitate (Figure 3A). The addition of palmitate to WT macrophages significantly increased OCR, suggestive of β-oxidation and oxidative phosphorylation (Figure 3B). Treatment of lipin-1 KO macrophages with palmitate failed to increase OCR (Figure 3C). There was a significant decrease in OCR in lipin-1 KO macrophages compared to WT macrophages with the addition of palmitate (Figure 3D). These data suggest that lipin-1 is critical for the breakdown and utilization of lipids during stimulation. A potential explanation for the inability to utilize fatty acids is due to a defect in uptake. To address if lipin-1 regulates fatty acid uptake, we treated WT and lipin-1 KO macrophages with BODIPY labeled palmitate and incubated for 1 hour at either 4°C or 37°C to differentiate cellular binding from lipid uptake. Cells were analyzed via flow cytometry to determine mean fluorescent intensity (MFI) as a marker for palmitic acid uptake. There was no difference in MFI between WT and lipin-1 KO macrophages (Figure 4A & 4B). Taken together these data suggest to us that the loss of lipin-1 from macrophages is not inhibiting lipid uptake, but rather lipid utilization.

**Figure 3:**
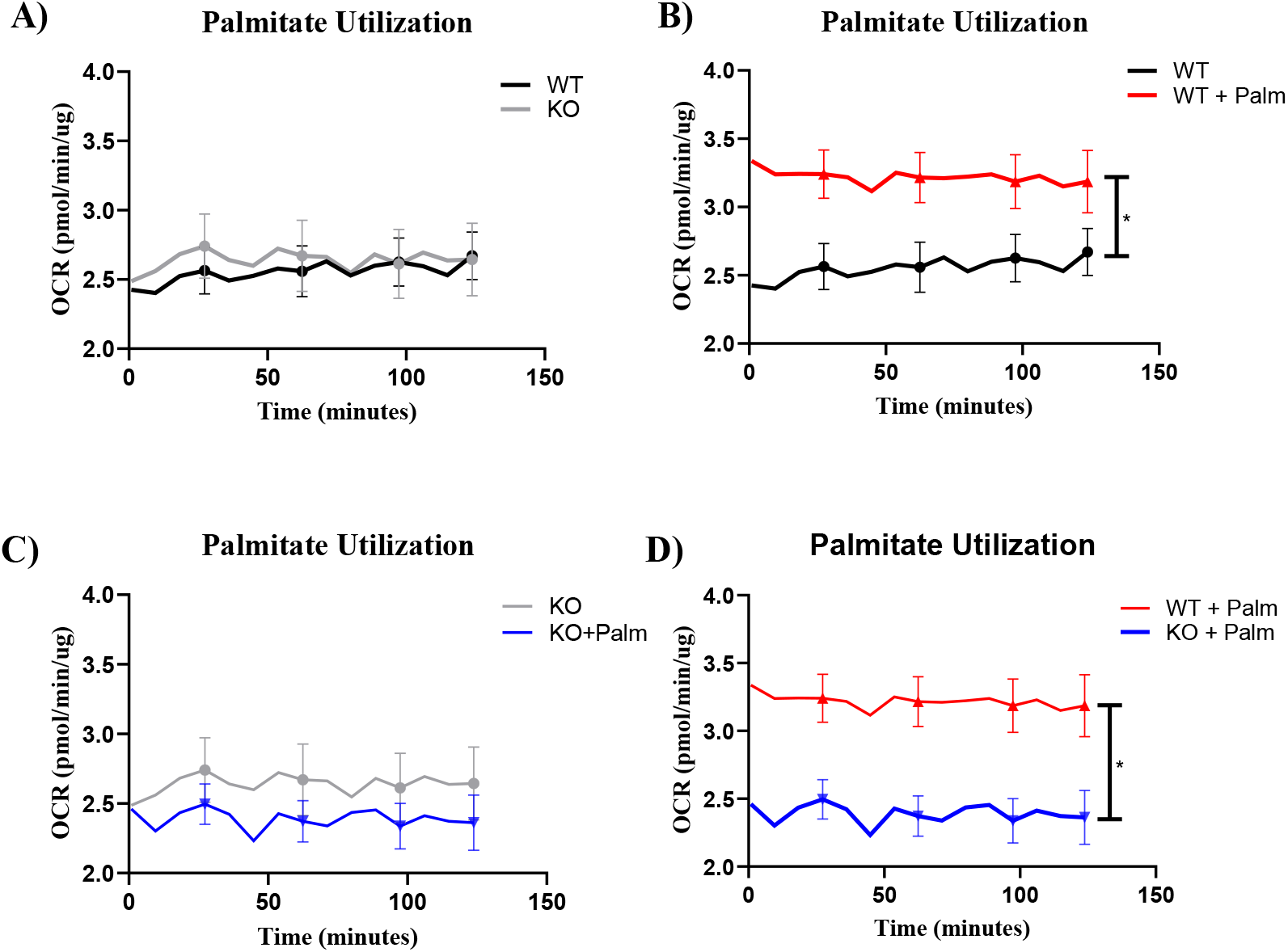
Lipin-1 is required for utilization of free fatty acids. BMDM isolated from WT and KO macrophages were treated with 40ng/mL for 1.5 hours. Immediately before assay, 1mM palmitate was added to requisite wells and OCR was analyzed via Seahorse extracellular flux assay. A) WT and KO BMDM’s not treated with palmitate. B) WT BMDM’s either treated or untreated with palmitate. C) KO BMDM’s either treated or untreated with palmitate. D) WT and KO BMDM’s treated with palmitate. Area under the curve was analyzed via one-way ANOVA. N=3. Stars represent p<0.05.

**Figure 4:**
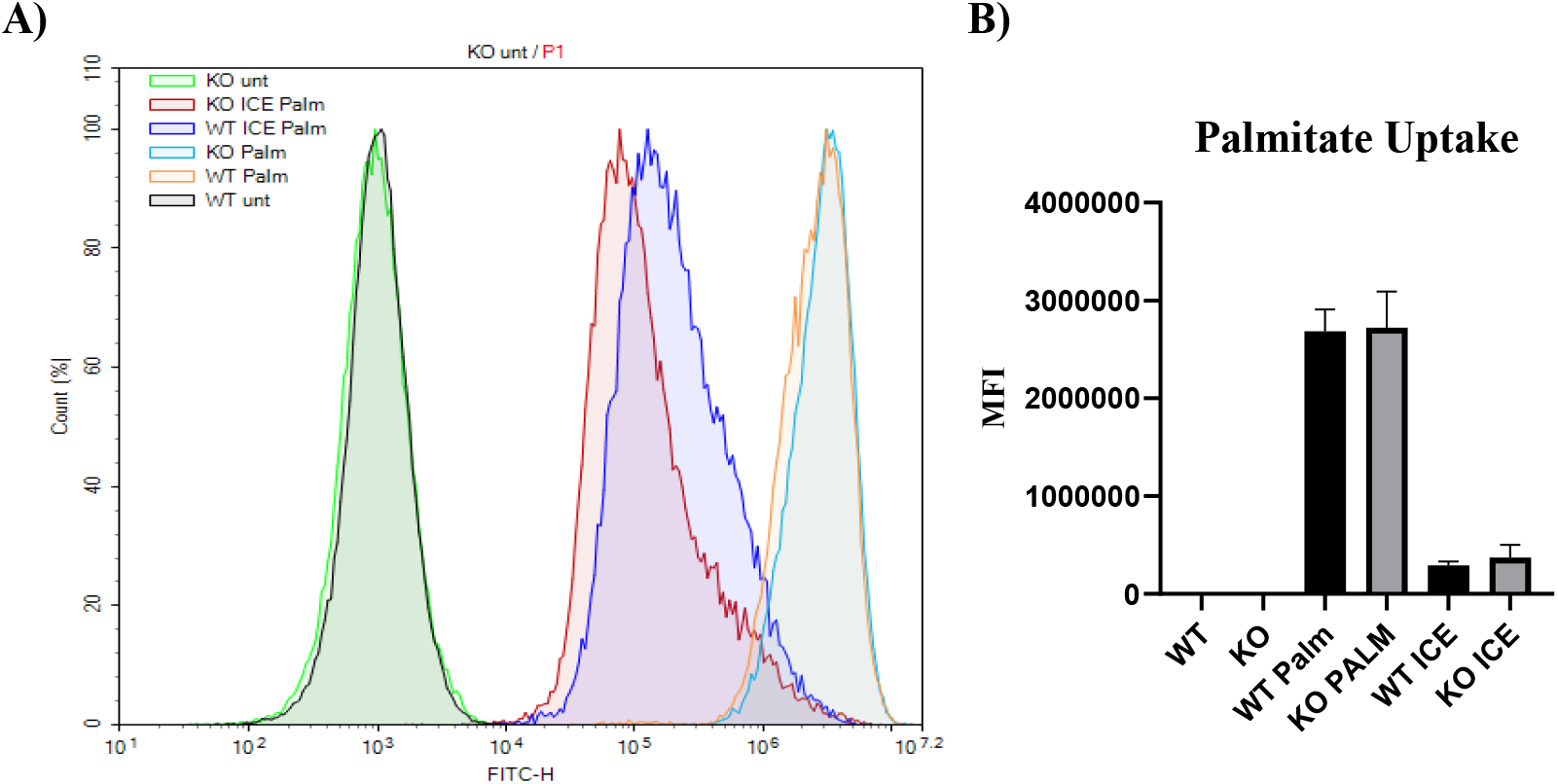
Lipin-1 does not regulate lipid uptake. BMDM isolated from WT and KO macrophages were treated with 100 μM BODIPY palmitate for 1 hour at either 37°C or at 4°C. BMDM’s were analyzed via flow cytometry to determine mean fluorescent intensity (MFI). A) A representative histogram of WT and KO BMDM’s treated with BODIPY palmitate. B) Compiled graph (N=3) of WT and KO BMDM’s treated with BODIPY palmitate. Data was analyzed via a student’s T-test.

### Lipin-1 regulates cellular metabolism

Our data suggests that lipin-1 is required for the breakdown and utilizing of lipid in response to IL-4 stimulation. Macrophages need to align individual metabolic pathways to effectively responds to stimuli. We wanted to understand the consequences of lipin-1 activity on immuno-metabolic responses in macrophages after stimulation with IL-4. WT and lipin-1 KO macrophages were stimulated with and without 40 ng/mL of IL-4 for 4 hours. Cell pellets were then sent to Creative Proteomics (Shirly, NY) for central carbon analysis to determine the abundance of intermediates in the major metabolic pathways such as glycolysis, TCA cycle, and pentose phosphate pathway. At baseline, lipin-1 deficient macrophages have an altered metabolism compared to WT macrophages. Lipin-1 KO BMDMs have multiple elevated glycolytic metabolites compared to WT BMDMs (Figure 5A & Supplemental figure 1). This data demonstrates that the loss of lipin-1 results in an increase in glycolysis. There is an overall decrease in basal TCA cycle intermediates with the exception of isocitrate, an isomer of citrate, in lipin-1 deficient BMDMs compared to WT BMDMs (Figure 5A & Supplemental figure 2). This suggests that there is a break in the TCA cycle of lipin-1 deficient BMDMs which leads to the accumulation of isocitrate in lipin-1 deficient BMDMs There is also an increase in the initial steps of the pentose phosphate pathway which is used to generate NADPH (Figure 5A).

**Figure 5:**
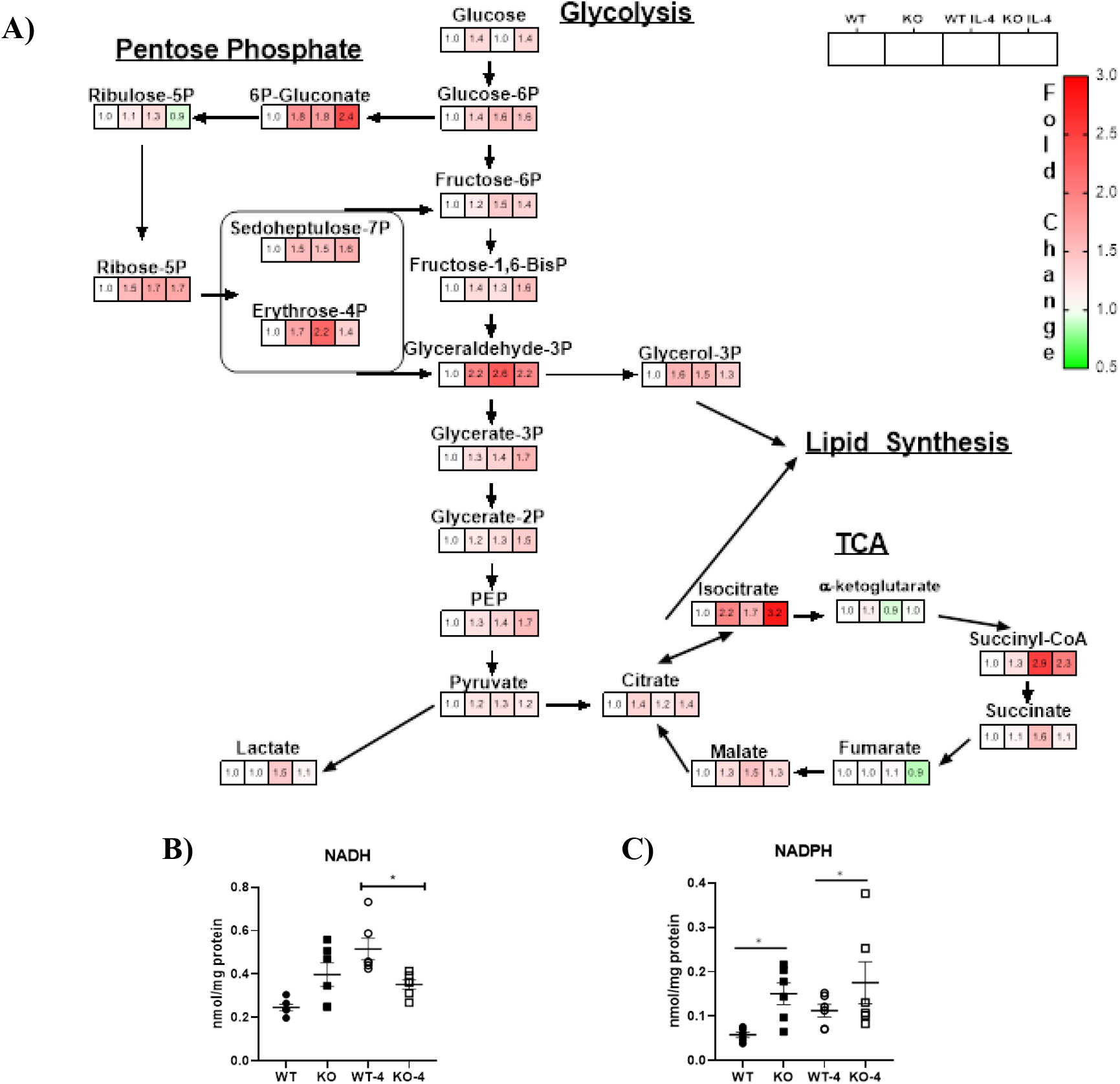
Lipin-1 regulates macrophage metabolism. Central carbon analysis of WT and KO BMDM’s treated with and without 40ng/mL IL-4 for 4 hours (A). Values represent fold change compared to WT control. Quantitative analysis of NADH (B) and NADPH (C) levels in WT and KO BMDM’s treated with and without 40ng/mL IL-4 for 4 hours. Student T-test was performed as statistical analysis. N=3. Stars represent p<0.05

Stimulation of WT BMDM with IL-4 resulted in a significant increase in many TCA cycle intermediates, consistent with a pro-resolving phenotype (Figure 5A & Supplemental figure 2). The alterations in metabolism in lipin-1 deficient BMDMs at baseline were exacerbated upon stimulation with IL-4. IL-4 stimulation of lipin-1 KO macrophage resulted in a significant increase in isocitrate, with an overall decrease in downstream metabolites (Figure 5A & Supplemental figure 2). These data suggest a break in the TCA cycle that is phenotypically similar to pro-inflammatory macrophages, which have decreased β-oxidation and oxidative metabolism[23]. The TCA cycle generates electron carriers (NADH) that are used by the electron transport chain to generate ATP. The loss of lipin-1 resulted in a significant decrease in NADH levels compared to WT macrophages (Figure 5B), further supporting a break in the TCA cycle of lipin-1 KO macrophages. Lipin-1 deficient macrophages have an increase in the initial metabolites of the pentose phosphate pathway, which is used to generate NADPH (Figure 5A). The increase in initial metabolites of the PPP is further supported by significantly elevated NADPH levels (Figure 5C). These results suggest that lipin-1 coordinates lipid catabolism to promote the metabolic phenotype of pro-resolving macrophages. Complete central carbon analysis can be found in (Supplemental table 1).

### Lipin-1 promotes efferocytosis

Macrophages mediate tissue homeostasis by the clearance of dead and dying cells by a process termed efferocytosis. Ingestion of an apoptotic cell results in an overabundance of cellular material (sugars, amino acids, and lipids) that need to be processed to allow the macrophage to continue efferocytosis[6]. Recent studies have demonstrated that β-oxidation of AC derived lipids is necessary for IL-10 production and subsequent wound healing in a mouse model of myocardial infarction [6]. Defects in efferocytosis lead to unresolved inflammation that perpetuates disease pathogenesis, as seen in atherosclerosis, obesity, diabetes, and autoimmunity [24–26]. We have demonstrated that mice lacking myeloid associated lipin-1 have increased necrotic cores within atherosclerotic plaques and have delayed wound closure [19, 20]. Clearance of dead cells within wounds is critical for effective wound closure, as well as reducing necrotic core formation [27, 28]. We re-examined data collected from [19] of the number of macrophages to the number of dead cells within wounds of lipin-1^m^KO mice and littermate controls. Although there was no difference in the total number of dead cells in the wound between lipin-1^m^KO mice and littermate controls, we observed a greater ratio of macrophages/dead cells within wounds of lipin-1^m^KO mice as compared to littermate controls. This may suggest that macrophages from lipin-1^m^KO mice may have a defect in efferocytosis (Figure 6A).

**Figure 6:**
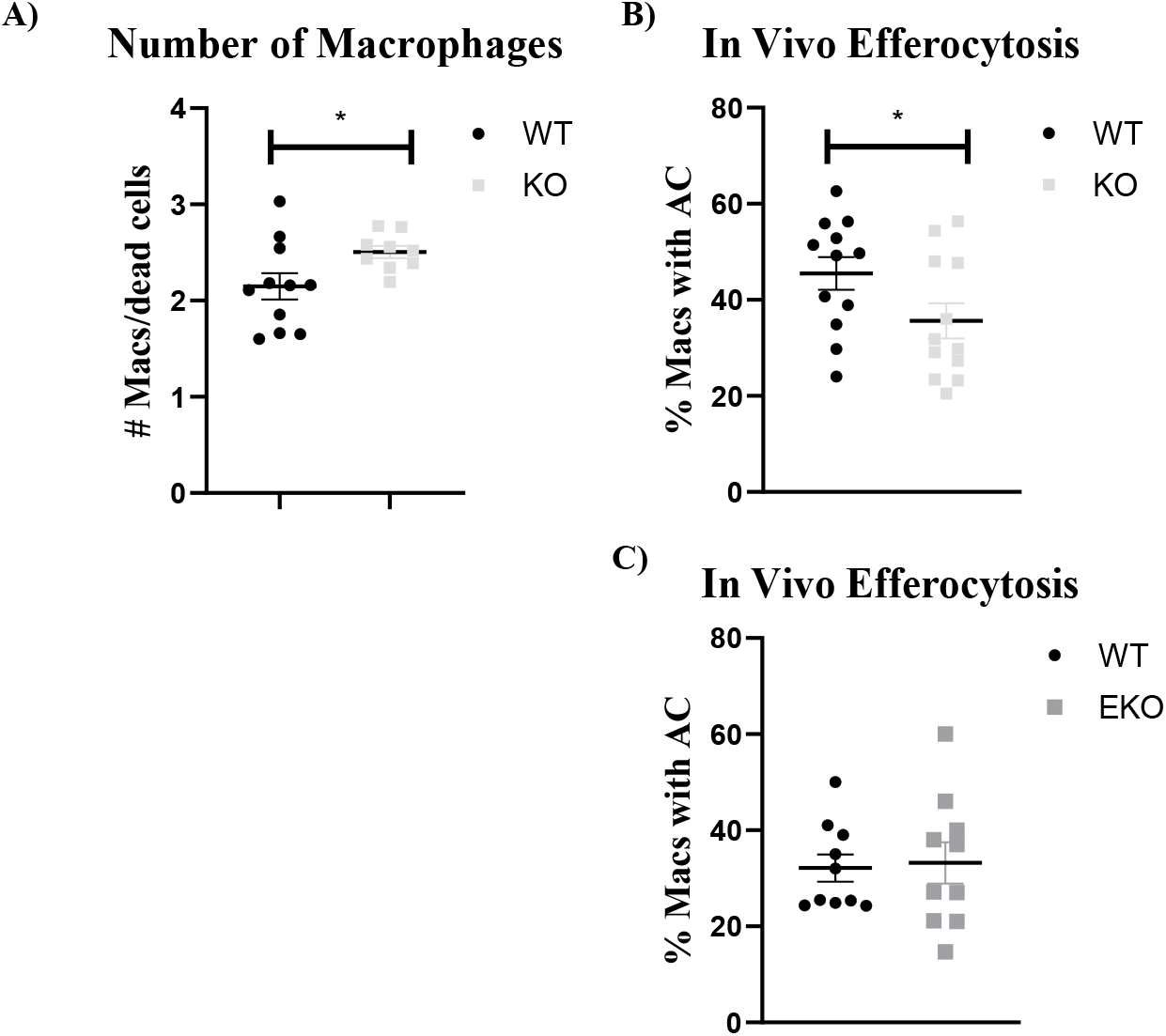
Lipin-1 contributes to efferocytosis. A) WT and KO mice subjected to an excisional wound healing model. Number of macrophages/dead cells were analyzed via flow cytometry. WT, KO (B), and EKO (C) mice were subjected to a zymosan model of peritonitis followed by peritoneal injection of PHK26 labeled AC. Percentage of macrophages with labeled AC were analyzed via flow cytometry. Students T-test was used to analyze data. N= 10-12. Stars indicate p<0.05

We decided to determine if lipin-1 is required for efferocytosis. To examine efferocytosis, we subjected lipin-1^mEnzy^KO, lipin-1^m^KO and littermate control mice to a zymosan model of inflammation. Briefly, mice were injected intraperitoneally with 0.1mg of zymosan. The inflammatory response was allowed to continue for 6 days to establish a pro-resolving milieu [29]. Labeled AC were injected into the peritoneal cavity for 45 minutes. Peritoneal lavages were collected and percentages of macrophages with labeled AC were quantified via flow cytometry. Lipin-1^m^KO mice had a significant reduction in the percentage of macrophages with AC compared to both WT and lipin-1^mEnzy^KO (Figure 6B & 6C). These data demonstrate that lipin-1 enzymatic activity is not involved in efferocytosis, and suggest the transcriptional coregulatory activity promotes efferocytosis in macrophages. Lipin-1 transcriptional coregulatory function regulates PPARs, which have been shown to increase efferocytic receptors such as MerTK [8, 30]. A reduction in efferocytosis receptors in lipin-1 KO macrophages could explain the defect in efferocytosis. To determine if the loss of lipin-1 results in an alteration in cell surface expression of MerTK, we stained peritoneal lavages for macrophages (CD11b and F4/80) and analyzed the presence of MerTK via flow cytometry. There was no significant difference in cell surface expression of MerTK in WT and lipin-1 KO macrophages (Supplemental figure 3), suggesting the defect in efferocytosis is not due to the lack of MerTK.

We have shown that lipin-1 is required for increased OCR in response to free fatty acids, but we wanted to know if lipin-1 regulates β-oxidation of AC derived lipids during efferocytosis. BMDM from WT, lipin-1^mEnzy^KO, and lipin-1^m^KO mice were either pretreated with or without etomoxir to prevent fatty acid transport into the mitochondria followed by the addition of AC to the macrophages at an AC/BMDM ratio of 4:1 then OCR was analyzed via Seahorse XFe analyzer (Agilent Technologies). WT and lipin-1 EKO macrophages significantly increased OCR in response to AC, with a significant reduction with pre-treatment of etomoxir (Figure 7A & 7B). This suggests that the increase in oxidative metabolism is due to the catabolism of AC derived lipids in both WT and EKO macrophages. However, lipin-1 KO macrophages failed to respond to AC compared to WT macrophages, with no significant difference in OCR with pretreatment of etomoxir (Figure 8A & 8B). These data suggest that the transcriptional coregulatory function of lipin-1 is required for β-oxidation of AC-derived lipids during efferocytosis, while the enzymatic activity of lipin-1 is not required.

**Figure 7:**
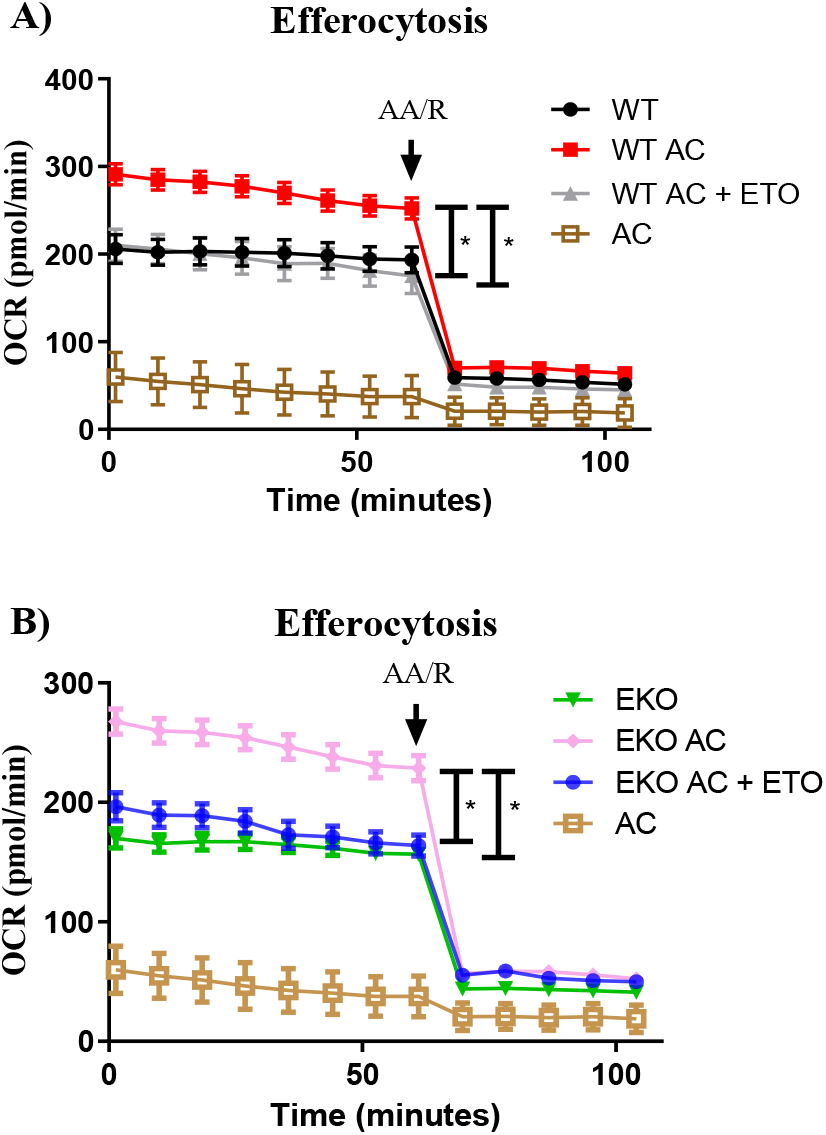
Lipin-1 enzymatic activity does not contribute to AC cell derived lipid utilization. Oxygen consumption rate of WT (A) and EKO (B) BMDM’s pretreated with and without 40 μM etomoxir for 15 minutes followed by the addition of AC at a 4:1 ratio. N=3. Stars indicate p<0.05

**Figure 8:**
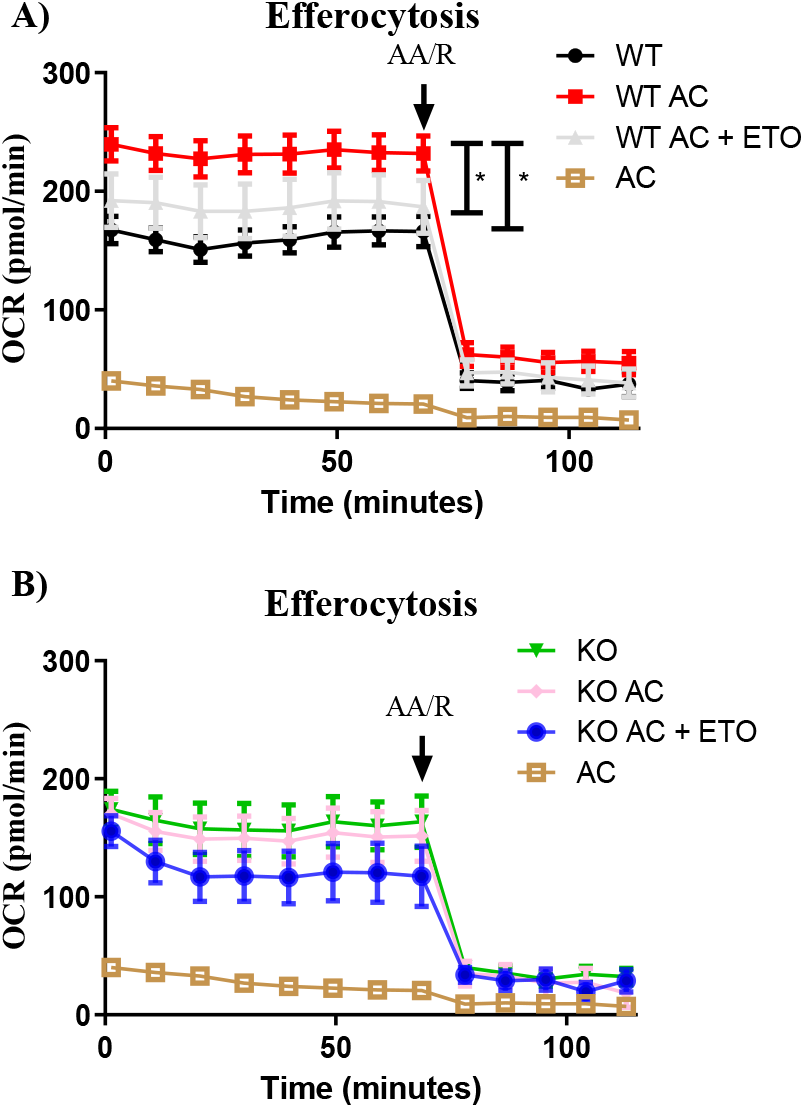
Lipin-1 transcriptional coregulator function is required for degradation of AC cell derived lipids. Oxygen consumption rate of WT (A) and KO (B) BMDM’s pretreated with and without 40 μM etomoxir for 15 minutes followed by the addition of AC at a 4:1 ratio. N=3. Stars indicate p<0.05

## DISCUSSION

Pro-resolving macrophages undergo drastic metabolic changes to effectively respond to stimuli. These pro-resolving macrophages significantly increase β-oxidation and oxidative phosphorylation, and perturbations in these pathways are deleterious to macrophages function[5]. We have previously demonstrated that lipin-1 contributes to macrophage polarization towards a pro-resolving phenotype, and atherosclerotic progression[16, 19, 20]. In both circumstances, lipid metabolism is crucial for appropriate macrophage function. Through the use of our Lipin-1^mEnzy^KO, lipin-1^m^KO, and littermate control mice, we provide evidence that lipin-1 transcriptional coregulator function is critical for appropriate lipid metabolism during IL-4 stimulation as well as during efferocytosis. We also provide evidence that alterations in lipid metabolism negatively impacts the pro-resolving functions of macrophages, which may contribute to impairment in disease resolution.

IL-4 stimulation as well as efferocytosis leads to activation of transcription factors such as STAT6 and PPARs to drive expression of genes involved in fatty acid transport and β-oxidation[30, 31]. Lipin-1 binds to and augments PPAR activity in a variety of tissue[8, 9]. The removal of exons 3 and 4 of lipin-1 in our lipin-1^mEnzy^KO mice results in truncated lipin-1 that lacks enzymatic activity but retains the ability of lipin-1 to bind to transcription factors such as PPARα and PPARγ[32]. Lipin-1^mEnzy^KO macrophages had equivalent levels of OCR in response to both IL-4 stimulation and AC. Removal of exon 7 from lipin-1 in our lipin-1^m^KO mice causes a missense protein leading to loss of lipin-1 and both activities[22]. BMDM from lipin-1^m^KO mice showed a significant reduction in OCR during stimulation of IL-4 and exogenous palmitate. There was an equivalent level of BODIPY palmitate uptake between WT and lipin-1 KO macrophages suggesting that the defect in β-oxidation is not due to the inability of the macrophages to take in the lipid. Further studies will investigate trafficking and utilization of the lipid within the macrophage. This suggests the transcriptional coregulator function of lipin-1 is responsible for regulating β-oxidation within macrophages.

Macrophage metabolism is directly connected to their polarization state and role during the inflammatory response[5]. Central carbon analysis of WT and lipin-1 KO macrophages untreated and treated with IL-4 revealed an overall increase in glycolysis in lipin-1 deficient macrophages. Interestingly, the loss of lipin-1 led to a significant decrease in lactate, with no alteration in pyruvate. Lactate can be shuttled into the mitochondria to be converted to pyruvate to fuel the TCA cycle [33]. Untreated and IL-4 treated lipin-1 KO macrophages have an accumulation of isocitrate, an isomer of citrate. Downstream TCA cycle metabolites after isocitrate were reduced in lipin-1 KO macrophages compared to WT macrophages. This is further supported by a decrease in NADH levels as well as a significant decrease in OCR in lipin-1 deficient macrophages. Isocitrate is an isomer of citrate which can be exported out of the mitochondria to be used in lipid synthetic pathways, similar to pro-inflammatory macrophages [34]. Lipin-1 deficient macrophages have an increase in the initial metabolites of the PPP, resulting in increased NADPH levels. NADPH is used by the enzymes of lipid synthetic pathways, as well as for redox potential with the cell[35]. Glycerol-3-phosphate is used as a backbone for glycerol lipid synthesis and is elevated in the lipin-1 KO macrophage (Supplemental 4). All together the loss of lipin-1 results in a metabolic profile that is supportive of lipid synthesis, rather than lipid catabolism. Interestingly, previous studies have shown that de novo lipid synthesis, such as ceramides, are detrimental to efferocytosis, supporting data shown in our current study [36]. The inability to undergo β-oxidation of exogenous lipid, together with the metabolic profile of lipin-1 KO macrophages may suggests that the incoming free fatty acids are being used for de novo lipid synthesis. Future studies will investigate how the loss of lipin-1 results in lipid synthesis.

Efferocytosis is a complex process that leads to the accumulation of macromolecules within the macrophage that need to be processed and cleared. Apoptotic cell derived arginine is metabolized leading to RAC1 activation and further actin polymerization to allow for uptake of multiple AC[37]. Lipid is a major component of apoptotic cells. Mitochondria are critical for the process of efferocytosis[6, 38]. Mitochondria must undergo fission to mobilize around the engulfed AC to allow for continuing efferocytosis[38]. This is presumably to accept incoming fatty acids for β-oxidation. Β-oxidation of AC derived lipids is required for anti-inflammatory cytokine production and wound healing in a mouse model of myocardial infarction[6]. These studies eloquently show the importance of the mitochondria to efferocytosis. Here we show that lipin-1 regulates lipid catabolism and we propose the defect in efferocytosis may be due to the inability to breakdown lipid. Future studies will investigate the mechanism in which lipin-1 regulates efferocytosis. We provide evidence that the transcriptional coregulatory function of lipin-1 is required for lipid catabolism both during treatment with IL-4, exogenous palmitate, and apoptotic cells. Our work highlights the importance of lipin-1 and lipid metabolism to macrophage function. Future work will provide a more mechanistic insight on the pathways that lipin-1 regulates to promote efferocytosis.

Our study demonstrates that lipin-1 is required for increased oxidative phosphorylation and lipid catabolism within pro-resolving macrophages. We believe that lipin-1 regulates the transcriptional profile needed for proper β-oxidation and oxidative metabolism. We also show that lipin-1 is required for efficient efferocytosis. Mice lacking myeloid associated lipin-1 have reduced AC uptake in a zymosan model of efferocytosis. Although unknown, we hypothesize that the defect in efferocytosis is due to an inability to breakdown AC derived lipid. Further studies will more mechanistically investigate how lipin-1 contributes to efferocytosis.

## ACKNOWLEDGMENTS

We would like to thank David Custis for his help with running samples related to flow cytometric experiments, and Gabrielle Gahn for determining genotypes of all mice.

