## Supplemental figures for "Lipin-1 regulates lipid catabolism in pro-resolving macrophages"

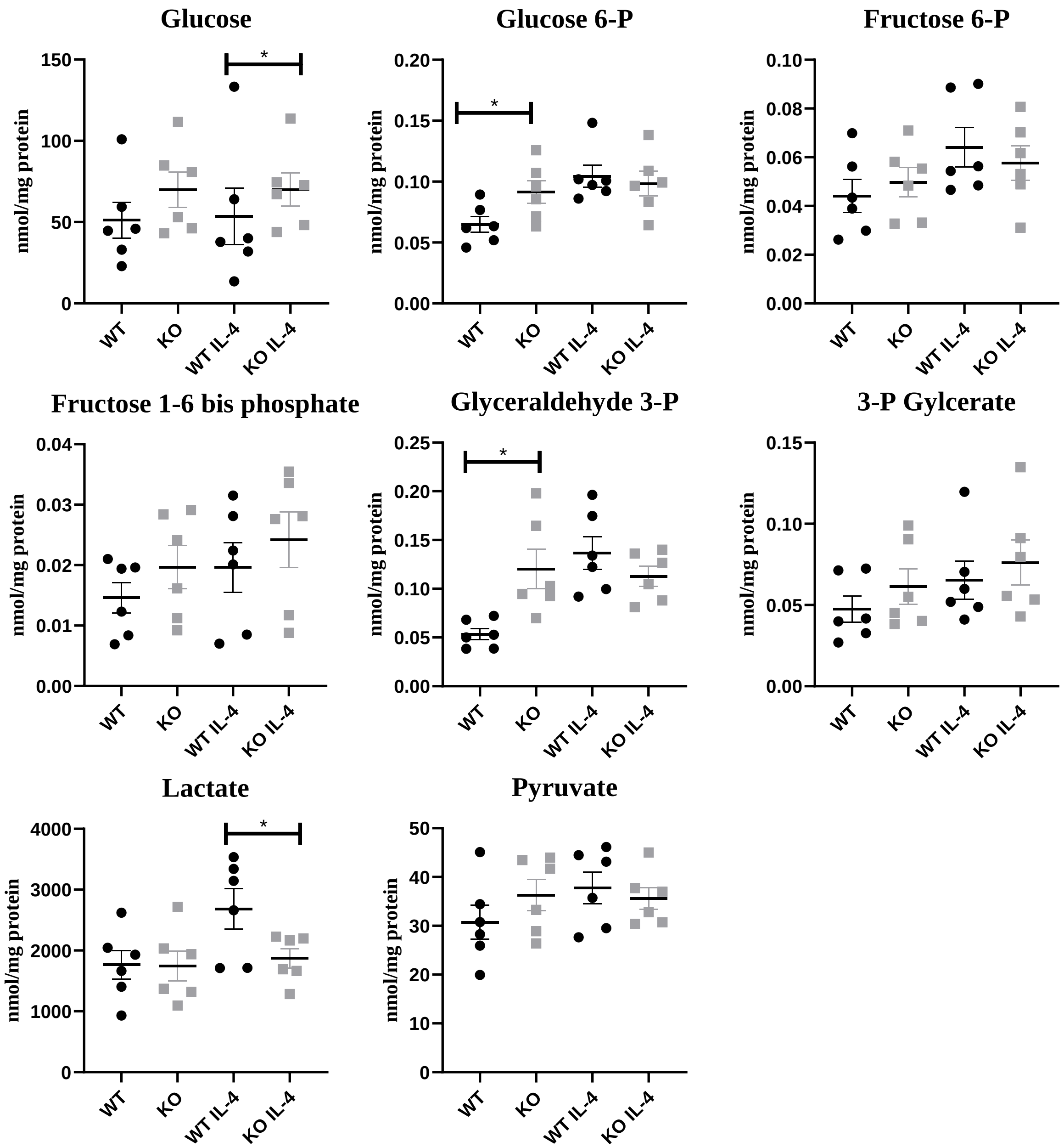


Supplemental figure 1: Lipin-1 coordinates metabolism with macrophages.

Central carbon analysis of WT and KO BMDM's treated with and without 40ng/mL IL-4 for 4 hours. Glycolytic intermediates. N=3. Stars indicate $p<0.05$

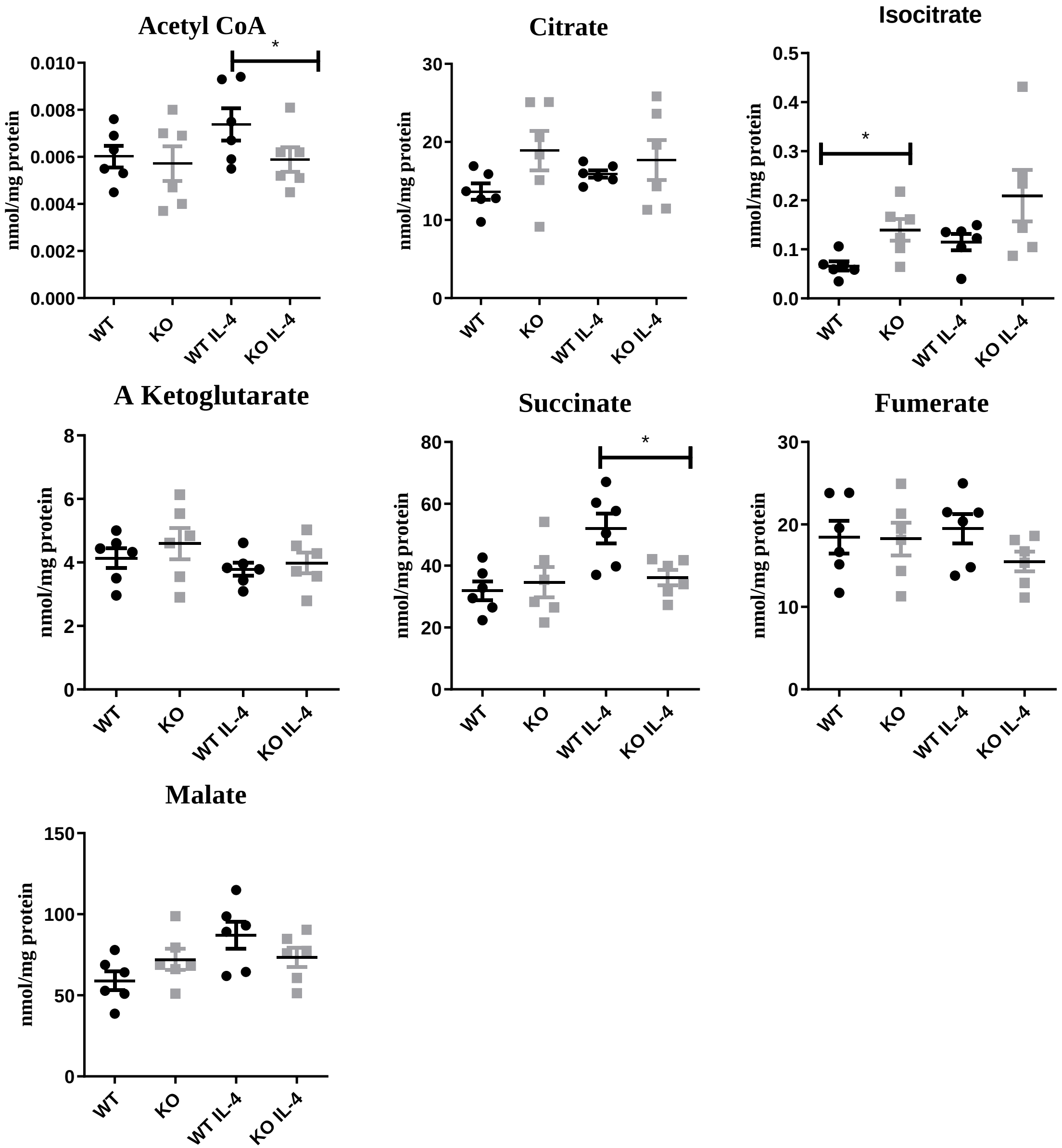


Supplemental figure 2: Lipin-1 coordinates metabolism with macrophages.

Central carbon analysis of WT and KO BMDM's treated with and without 40ng/mL IL-4 for 4 hours. TCA cycle intermediates. N=3. Stars indicate $p<0.05$

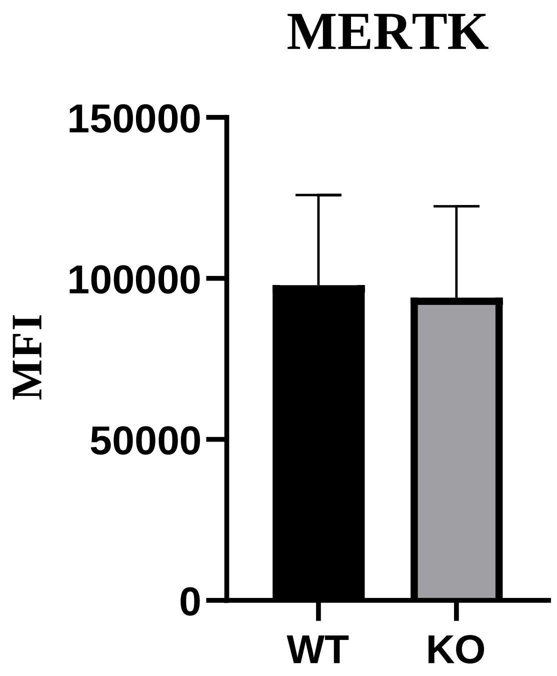


Supplemental figure 3: Lipin-1 does not regulate MerTK cell surface expression.

WT and KO mice were subjected to a zymosan model of peritonitis. Flow cytometry analysis of macrophage MerTK was performed on peritoneal lavages. N=2.

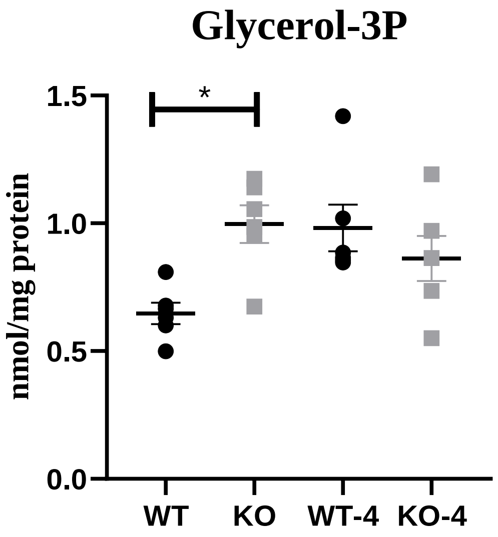


Supplemental figure 4: Lipin-1 deficiency leads to an increase in lipid synthetic intermediates.

Central carbon analysis of WT and KO BMDM's treated with and without 40ng/mL IL-4 for 4 hours. Quantitative analysis of glycerol-3 phosphate levels. N=3. Stars indicate $p<0.05$
